# What threatens Brazilian endangered species and how they are Red-Listed

**DOI:** 10.1101/711242

**Authors:** Enrico Bernard, Fernanda Silva de Barros, Vitória Edna Fernandes Felix, Thomas E. Lacher

## Abstract

Brazil is a continental-sized megadiverse country with high rates of habitat loss and degradation. Part of the Brazilian biodiversity – including endemic species – is recognized as threatened. By following the IUCN standards, we review the classification of all the 1172 endangered species in Brazil, analyzing differences among categories and groups. Based on a subsample of all 464 terrestrial vertebrates we identified 1036 records of threats affecting them. Criterion B was the most used (56% overall; 70% for CR species; 75% for EN), mainly related to reductions in their habitat area, extent and/or quality due to deforestation. Data on population declines (criterion A), number of reproductive individuals (criterion C), and population sizes (criterion D) are available for only a small fraction of the Brazilian fauna. Criterion E (probability of extinction in the wild) was used for only one species. Birds and mammals had the highest diversity of used criteria, while marine fish the lowest (90% related to declining populations). Two out of three of the 464 vertebrate species analyzed were negatively impacted by agribusiness. Other major threats are hunting, urban sprawl, rural settlements, and the construction of hydroelectric dams. Birds and mammals experience more co-occurrence of threats. Some threats are clearly underestimated in Brazil: climate change was indicated for only 2% species analyzed, but included no birds or amphibians. The main threats identified are linked to the patterns of economic development in Brazil and the current political and economic context points to a worrisome conservation scenario in the near future.

## Abstract

Resumo

O Brasil é um país continental megadiverso com altas taxas de perda e degradação de habitats. Parte da biodiversidade brasileira – incluindo espécies endêmicas – está ameaçada. Seguindo padrões da IUCN, aqui detalhamos a classificação de todas as 1172 espécies ameaçadas de extinção no Brasil, analisando as diferenças entre categorias e grupos animais. Baseado em uma amostra de todas as 464 espécies de vertebrados terrestres ameaçados identificamos 1036 registros de ameaças sobre elas. O Critério B foi o mais usado (56% no geral; 70% para espécies CR; 75% para EN), principalmente em função de reduções na área, extensão e/ou qualidade dos habitats em função de desmatamento. Dados sobre declínios populacionais (critério A), número de indivíduos reprodutivos (critério C), e tamanho populacional (critério D) existem para apenas uma pequena fração da fauna brasileira. O Critério E (probabilidade de extinção na natureza) foi usado para apenas uma espécie. Aves e mamíferos têm a mais alta diversidade de critérios usados, enquanto peixes marinhos a menor (90% relacionados com declínios populacionais). Duas de cada três das 464 espécies de vertebrados analisadas são negativamente afetadas pelo agronegócio. Outras ameaças incluem a caça, expansão urbana, assentamentos rurais, e a construção de hidrelétricas. Aves e mamíferos experimentam a maior co-ocorrência de ameaças. Algumas ameaças estão claramente subestimadas no Brasil: mudanças climáticas foram indicadas apenas para 2% das espécies analisadas, mesmo assim para nenhuma ave ou anfíbio. As principais ameaças identificadas estão ligadas aos padrões de desenvolvimento econômico no Brasil e os contextos político e econômico atuais apontam para um cenário conservacionista preocupante no futuro próximo.

## Introduction

Drivers of environmental changes are increasing globally, pushing biodiversity loss at unprecedented rates in almost all ecosystems on the planet (e.g. Ceballos et al. 2015). The situation is more severe in tropical regions, with a complex combination of high species richness, increasing human populations, and high rates of natural habitat loss. But even in the Tropics the situation is heterogeneous, more worrisome in some countries. This is the case of Brazil, a country with some world records of biodiversity – including thousands of endemic species – but also very high rates of habitat loss and degradation and a pessimistic political scenario (e.g. Abessa et al. 2019; Gonzales, 2019; Phillips 2019).

In December 2014, after a hiatus of more than a decade, the Brazilian Minister of Environment updated the official list of endangered species in Brazil (MMA, 2014). Between 2010 and 2014, more than 1300 specialists evaluated 12,556 species in 73 workshops and four validation meetings (ICMBio, 2015a, 2015b). All known species of birds, mammals, amphibians and reptiles in Brazil were evaluated, plus 4507 fish and 3332 invertebrate species, and 1172 species were officially declared endangered in the country. Overall, that evaluation effort was more than 10 times larger (more than 20 times for marine and freshwater fish) than the previous one, published in 2003 (ICMBio, 2015b).

The list published in 2014 followed all the protocols set by the International Union for Conservation of Nature (IUCN), which adopt five quantitative criteria and sub criteria: A. Declining population (past, present and/or projected); B. Geographic range size, and fragmentation, decline or fluctuations; C. Small population size and fragmentation, decline, or fluctuations; D. Very small population or very restricted distribution; E. Quantitative analysis of extinction risk (e.g. Population Viability Analysis) (IUCN 2012; Supplementary Table 1).

Identifying threatened species is important, for example, for conservation prioritization purposes, regulation of trade in wildlife products, or for the legal protection of those species. However, diagnosing the threats experienced by a taxon or a group of taxa is critical to understand the risks they experience and, most importantly, to devise strategies to reverse their negative conservation scenarios. IUCN also adopts a unified classification system of threats (Salafsky et al., 2008), useful for conservation strategies because it defines and classifies threats in a standardized way and can be universally applied in different countries and contexts. Such classification has been used in recent research to determine the major threats to biodiversity globally (Maxwell et al., 2016) and regionally in Australia (Allek et al., 2018). The adoption of such internationally-applied criteria allows the scientific community, conservationists and decision-makers to better analyze and compare why species are threatened and how they were classified.

Here we provide a quali-quantitative analysis on how the Brazilian endangered species are being Red-Listed and what threatens them. We first identified which were the most-used IUCN criteria and sub criteria to classify all the 1172 endangered species in Brazil, and analyzed how they differ among categories and animal groups. Later, using a subsample of all endangered vertebrates (464 species of birds, mammals, reptiles and amphibians) we investigated in detail the threats those species face.

## Methods

We considered all 1173 species present in the List of Brazilian Fauna Species Threatened with Extinction (MMA, 2014). Between 2010 and 2014, 12,256 taxa of the Brazilian fauna were evaluated, including all vertebrates described for the country, for a total of 732 mammals, 1980 birds, 732 reptiles, 973 amphibians and 4507 fish (3,131 freshwater, including 17 rays, and 1,376 marine), plus 3332 invertebrates, including crustaceans, mollusks, insects, porifera, myriapods, among others (ICMBio, 2018). The Instituto Chico Mendes de Conservação da Biodiversidade (ICMBio) – the federal authority responsible for the evaluation process – carried out 73 assessments and four validation workshops. ICMBio worked formally together with IUCN for assessment standardizations and validations and the final list of threatened species was published in December 2014, containing 110 mammals, 234 birds, 80 reptiles, 41 amphibians, 353 bony fish (310 freshwater and 43 marine), 55 cartilaginous fish (54 marine and 1 freshwater), 1 hagfish and 299 invertebrates (MMA, 2014). In total, 448 species were classified as Vulnerable (VU), 406 as Endangered (EN), 318 as Critically Endangered (CR) and 1 as Extinct in the Wild (EW) (Supplementary Table 1).

For a detailed analysis of threats, a sub-sample containing all 464 species of threatened vertebrates (all terrestrial tetrapods in the assessment) were considered: 233 birds, 110 mammals, 80 reptiles and 41 amphibians, distributed in 37 orders and 114 families (Supplementary Fig. 1). For birds, the Alagoas curassow *Pauxi mitu* was not considered since it is classified as EW. In this sub sample, 82 species were CR, 176 EN, and 206 VU (Supplementary Fig. 2). For each of them, information about threats they experience were taken from official sources of the Ministry of the Environment, such as the available species’ National Action Plans and/or from the information provided during their assessment and evaluation process. Any references to threats found were compiled and tabulated in spreadsheets for each of the species, according to their biological group and threat category (Supplementary Table 2).

For 29 of those 464 species it was not possible to identify any related threat due to several factors: past or current reduction of the population by unknown causes; species without records of sightings for many years; decline in the number of mature individuals due to unknown causes; inference that the species is possibly extinct; taxonomic uncertainty; highly endemic species; and species with very restricted occurrence areas and/or areas of occupancy. However, in the 2014 list, no supporting information was found for the lizard *Liolaemus occiptalis* (CR), and therefore data from the previous 2003 list were used. For the lizard *Tropidurus psammonastes* (EN) no information was found from either the current list or the 2003 list. This species is classified as Data Deficient (DD) on IUCN Red List (IUCN, 2018).

The identified threats were classified into 11 drivers and 40 sub-drivers as proposed by Salafsky et al. (2008)(Supplementary Table 2). The driver Geological Events (Sub-drivers *Volcanoes, Earthquakes/tsunamis, Avalanches/landslides*) was not considered, due to the irrelevance of such activities to the Brazilian fauna. Five modifications were necessary in the categories proposed by Salafsky et al. (2008), and all were made with the intention of increasing the clarity and adequacy of the categories originally proposed (see Methods in the Supplementary information).

## Criteria and sub criteria used

Among the 1172 species analyzed, 1050 species (97%) were classified based on a single criterion, 113 species were classified based on two, and nine species based on three criteria, resulting in a total of 1,303 assigned criteria. Overall, criterion B (geographic range size) was the most widely used (56%), followed by criterion A (declining populations – 24%), criterion D (very small or restricted populations – 11%), and criterion C (small population size and estimated continuing decline in the number of mature individuals – 8%) (Fig. 1; Table 1). Criterion E, which estimates the probability of extinction in the wild, was used for only one species, the maned-wolf *Chrysocyon brachyurus,* however in combination with another (A3). When endangered status is considered, criterion B was the most common for categories CR and EN (70% and 75% of the classifications, respectively), but the distribution of categories was more uniform for VU: 35% for criterion A, 27% for B, 26% for D, and 10% for C

**FIG. 1.**
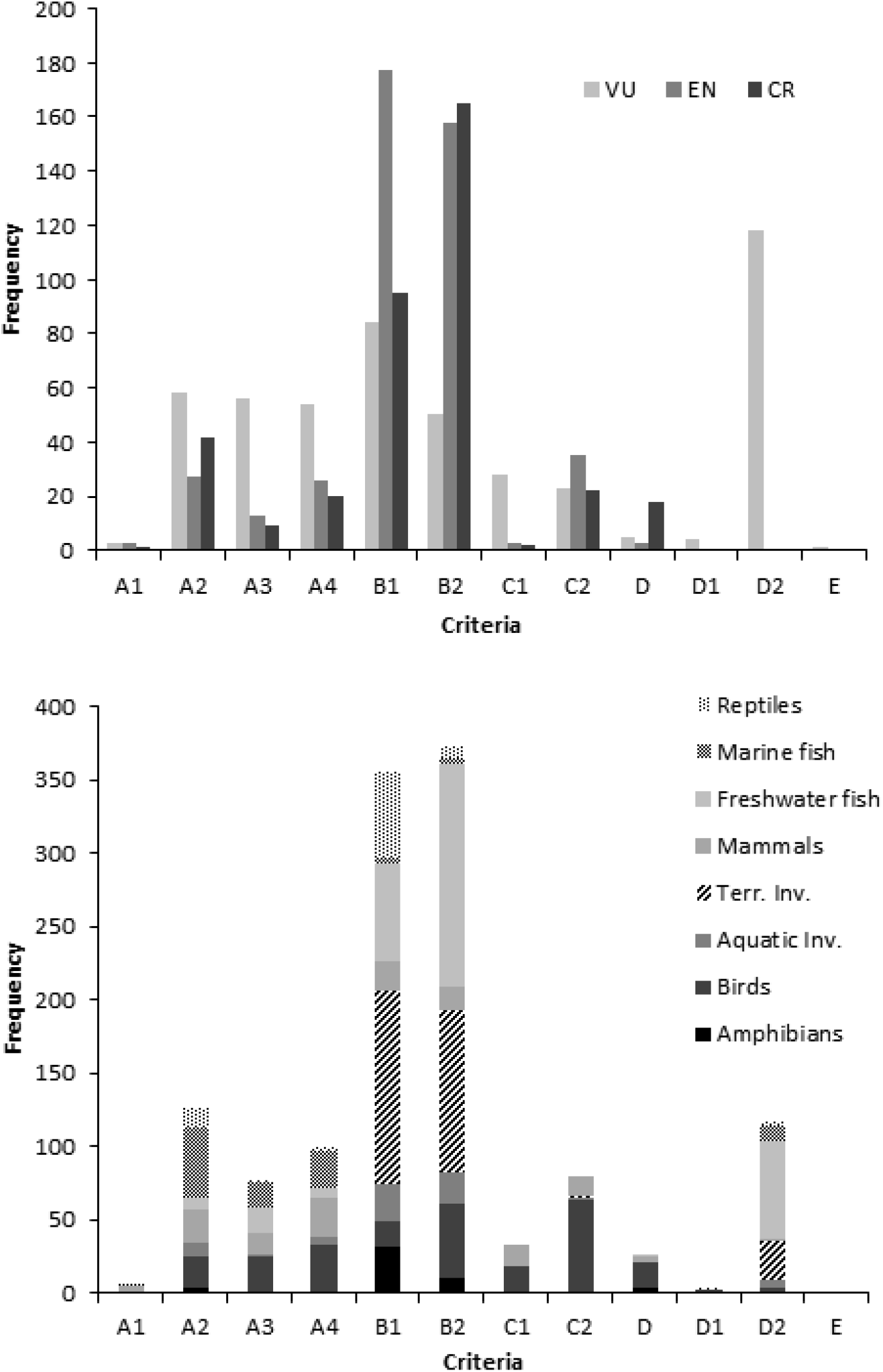
Distribution of IUCN Red List criteria for 1172 threatened species in Brazils according to threatening status (top) and animal group (bottom). VU = Vulnerable, EN = Endangered, CR = Critically Endangered.

**Table 1.**
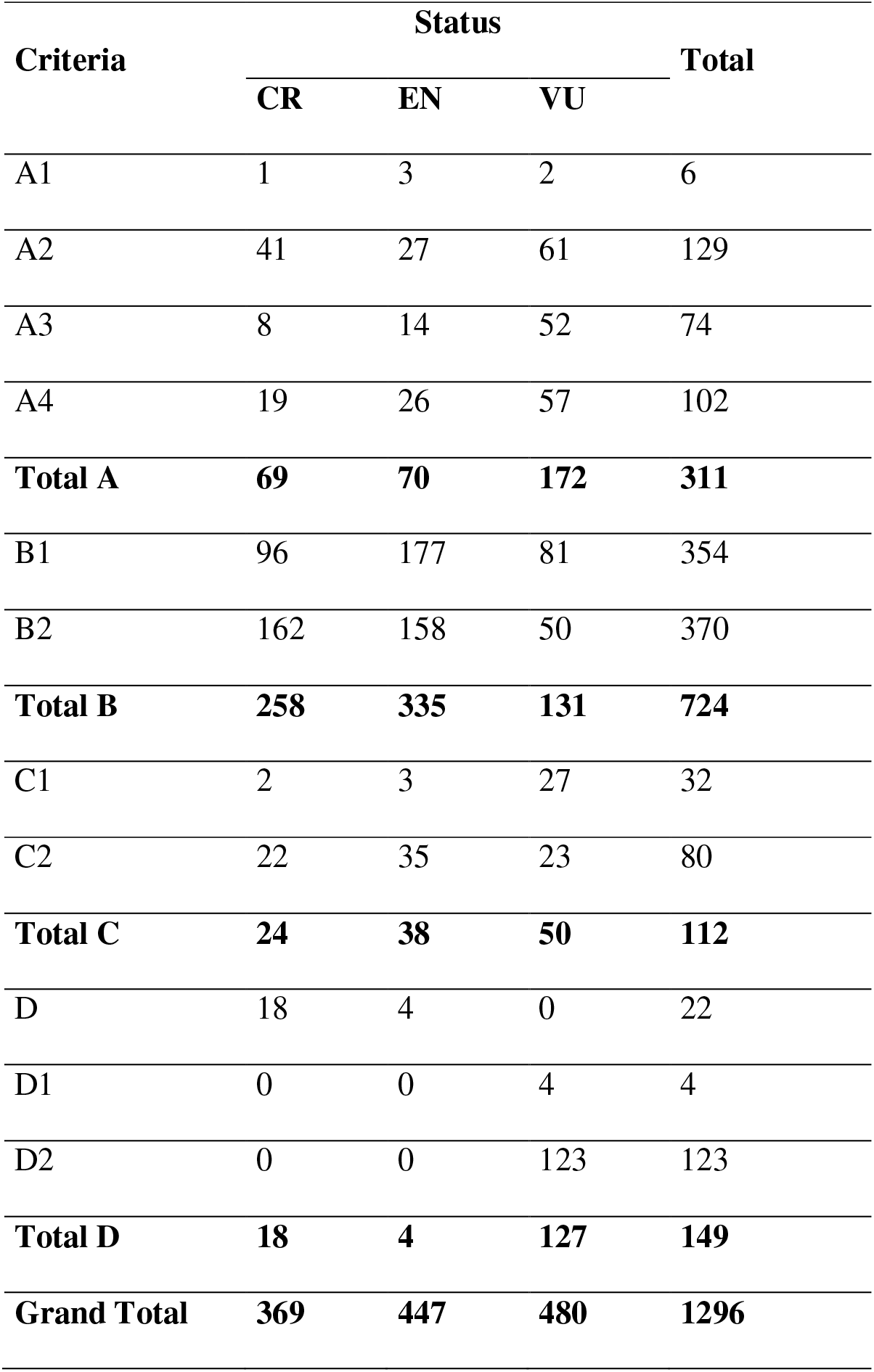
IUCN Red List criteria adopted and threat status for 1172 species officially listed as threatened in Brazil in 2014. Some species were listed under two or more criteria, so the total number of criteria used is higher than the number of species classified. CR = Critically Endangered, EN = Endangered, VU = Vulnerable.

| Criteria | Status |  |  | Total |
| --- | --- | --- | --- | --- |
|  | CR | EN | VU |  |
| A1 | 1 | 3 | 2 | 6 |
| A2 | 41 | 27 | 61 | 129 |
| A3 | 8 | 14 | 52 | 74 |
| A4 | 19 | 26 | 57 | 102 |
| **Total A** | **69** | **70** | **172** | **311** |
| B1 | 96 | 177 | 81 | 354 |
| B2 | 162 | 158 | 50 | 370 |
| **Total B** | **258** | **335** | **131** | **724** |
| C1 | 2 | 3 | 27 | 32 |
| C2 | 22 | 35 | 23 | 80 |
| **Total C** | **24** | **38** | **50** | **112** |
| D | 18 | 4 | 0 | 22 |
| D1 | 0 | 0 | 4 | 4 |
| D2 | 0 | 0 | 123 | 123 |
| **Total D** | **18** | **4** | **127** | **149** |
| **Grand Total** | **369** | **447** | **480** | **1296** |

Dozens of sub criteria combinations were used but overall the top-3, accounting for 66%, were B2 (area of occupancy severely fragmented or low number of locations – 29%), B1 (small extent of occurrence – 27%) and A2 (population reduction observed, estimated, inferred, or suspected in the past – 10%)(Fig. 1; Table 1). The high proportion of the use of criterion B2 was inflated by freshwater fish (47% of the 319 criteria for the group) and terrestrial invertebrates (41% of the 270 criteria for the group). For different categories of threat, the top-3 were B2 for CR species (44% out of 374 species), D2 for EN species (35% out of 445 species), and D2 for VU species (24% out of 484 species) (Fig. 1; Table 1).

Birds and mammals were the two taxonomic groups with the highest diversity of criteria used, but C2 and A4 were the most used, respectively (Fig. 2). For the other groups, criteria usage was more restricted: only four for amphibians and terrestrial invertebrates; and seven for the other groups. Terrestrial invertebrates and marine fish were the groups with the lowest diversity in usage, with 90% of the criteria being either B1 or B2 for the first, and 83% A1-A4 for the second.

**FIG. 2.**
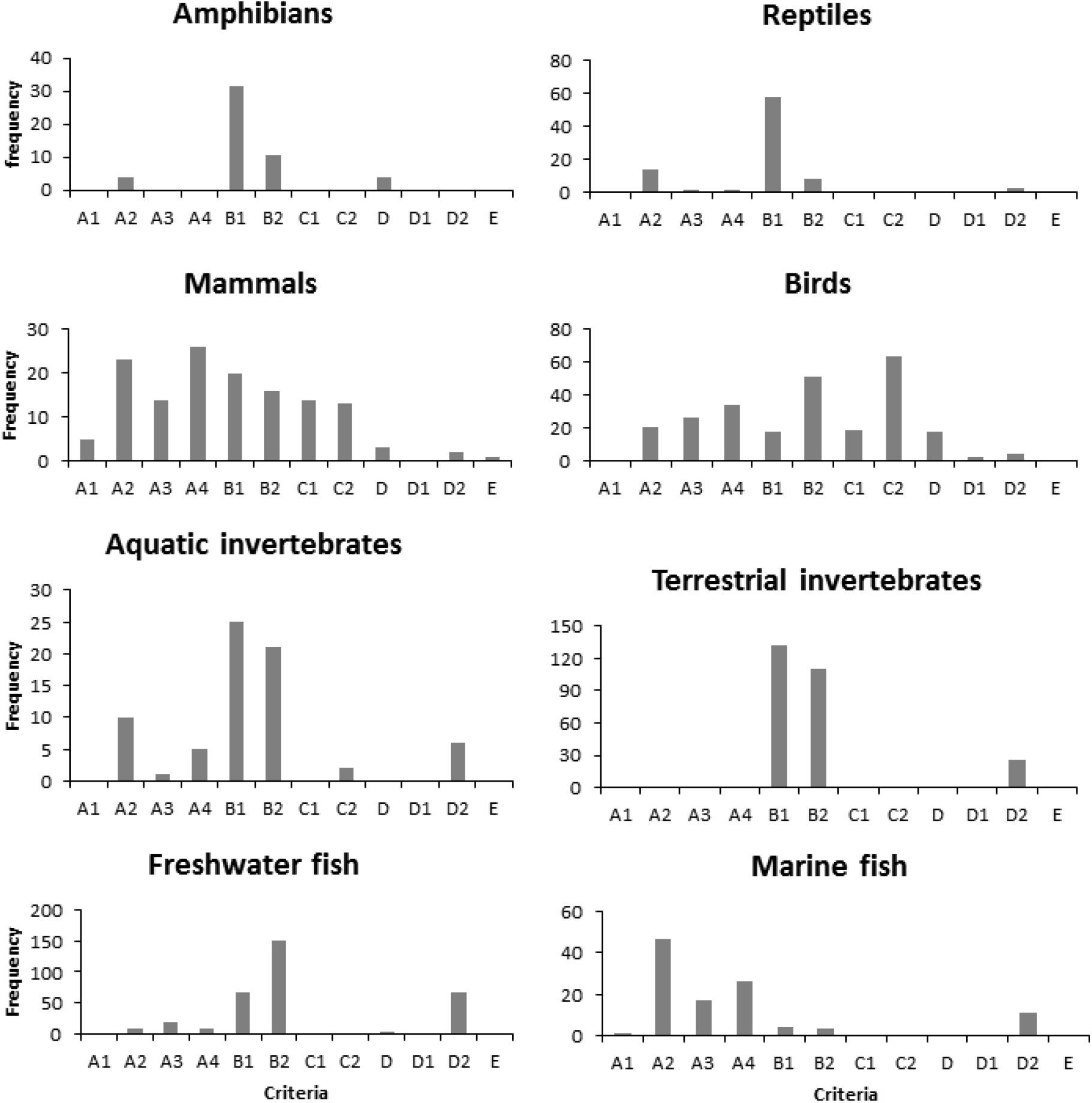
Distribution of IUCN Red List criteria for 1172 threatened species in Brazils according to animal group.

## Drivers, sub drivers and taxonomic biases

We identified 1036 records of threats for the 464 vertebrate species analyzed (Supplementary Table 2). The number of threats per species varied from 1 up to 10, with an average of 2.2 threats/species; 168 species had a single threat recorded, and 58% of the species had two or more simultaneously. The hooded capuchin monkey *Sapajus cay* (VU), with 10 records, the puma *Puma concolor* (VU) and the Coimbra-Filho’s titi monkey *Callicebus coimbrai* (EN), both with nine records, were the species with the highest number of threats identified.

The three most frequent drivers were *Agriculture and Aquaculture* (affecting 309 species), *Natural System Modification* (132 spp.), and *Overexploitation* (128 spp.), while the least frequent were *Pollution* (30 spp.), *Climate Change* and *Severe Weather* (10 spp.), and *Human Intrusions and Disturbance* (4 spp.) (Figs. 3 and S3). When species’ threat category is considered, *Agriculture and Aquaculture* affected most of the CR species (39 spp.), EN (120 spp.) and VU (150 spp.) (Fig. 4).

**FIG. 3.**
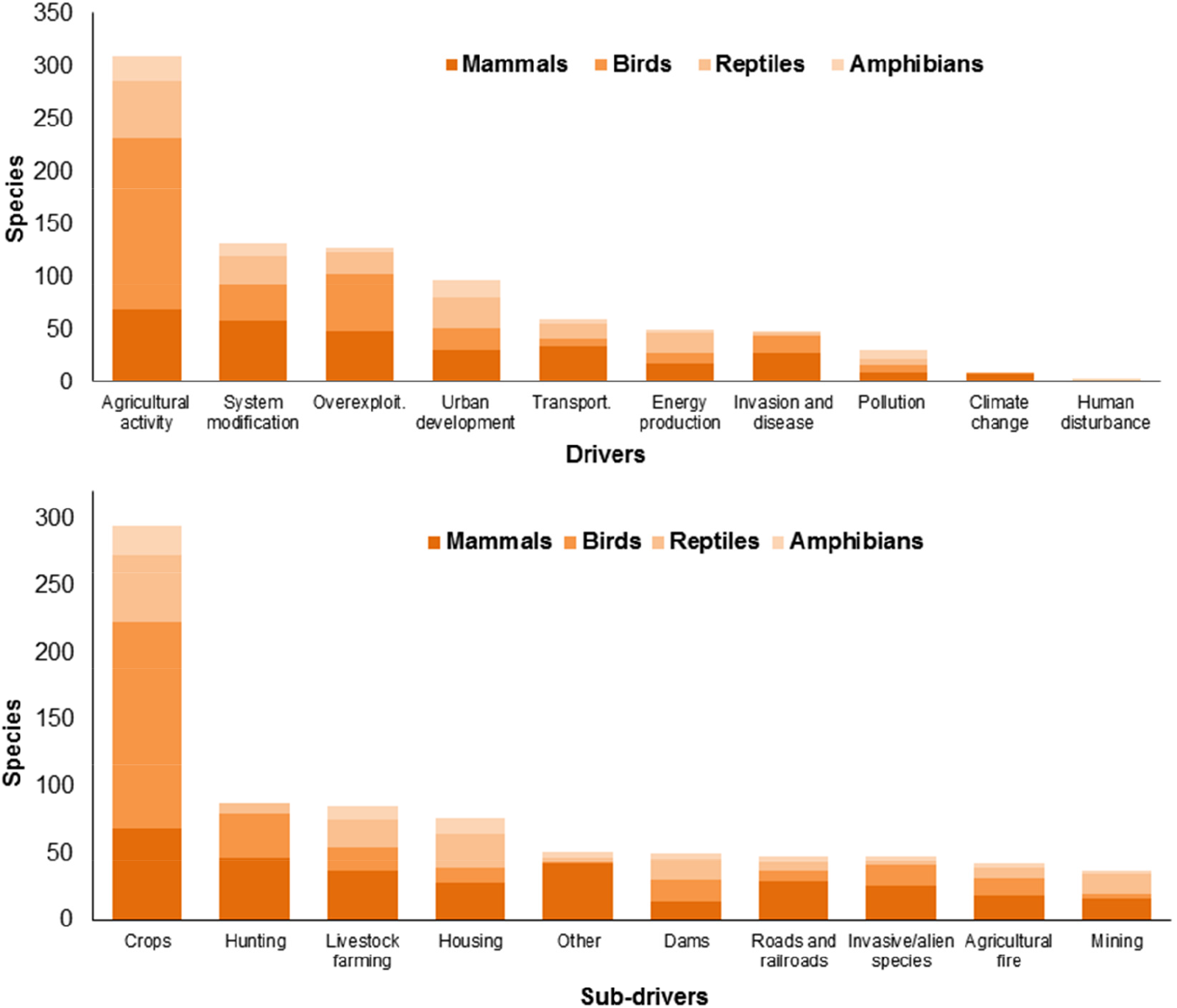
Top-10 threatening drivers (top) and sub-drivers (bottom) which affect the conservation of 464 threatened species mammals, birds, reptiles and amphibians in Brazil. Drivers were classified according to Salafsky et al. (2008).

**FIG. 4.**
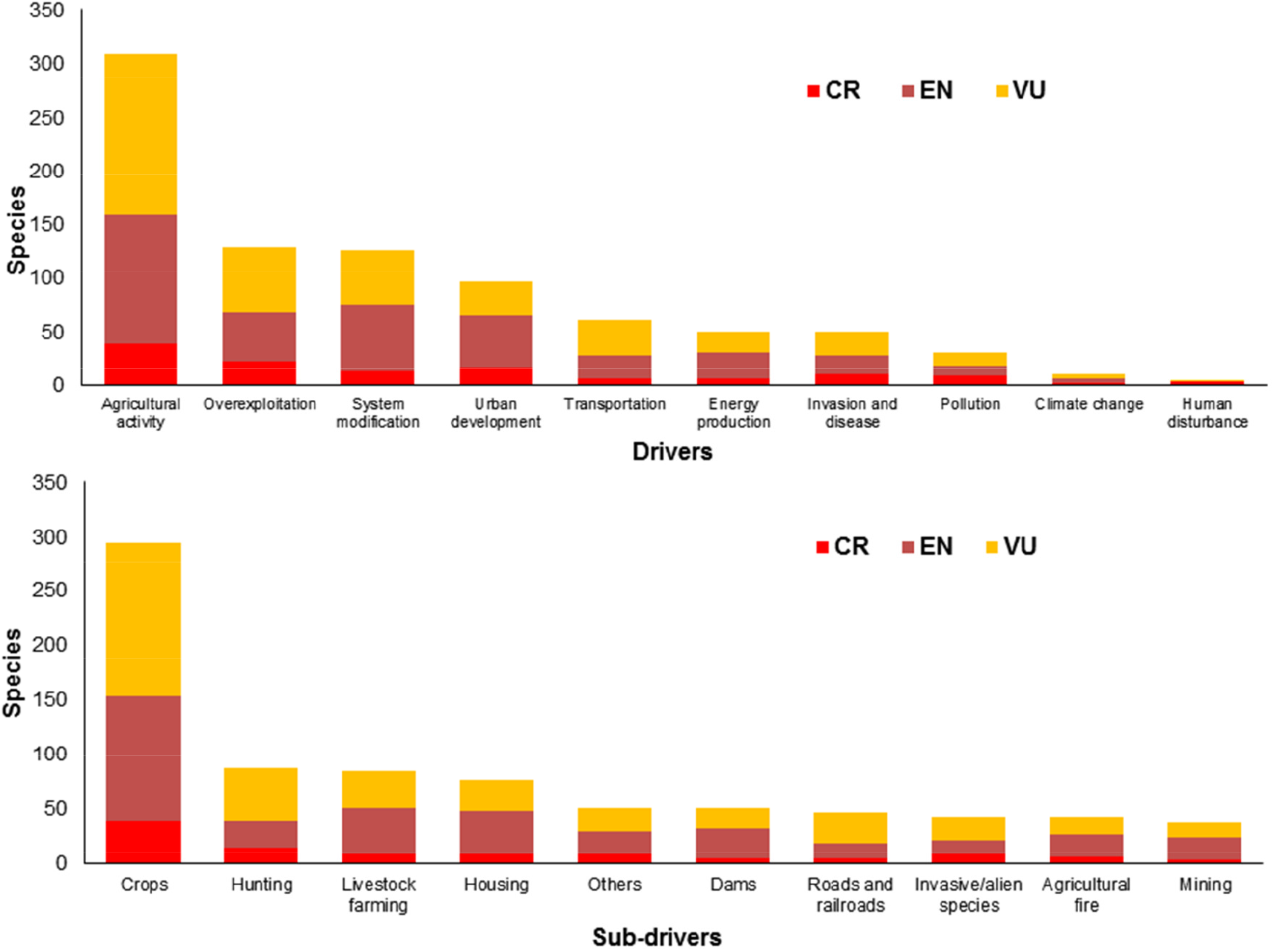
Top-10 threatening drivers (top) and sub-drivers (bottom) which affect the conservation of 464 threatened species of terrestrial vertebrates in Brazil. Drivers were classified according to Salafsky et al. (2008). CR = Critically Endangered, EN = Endangered, VU = Vulnerable.

The five most frequent sub-drivers were *Cropping* (for 295 species), *Hunting* (87 spp.), *Livestock farming* (85 spp.), *Housing* (76 spp.), and *Others* (51 spp.) (Figs. 3 and Supplementary 4). *Cropping* and *Livestock farming* were always among the three most frequent sub-drivers for all animal groups (Supplementary Fig. 5). The sub-driver *Agricultural fire* was recorded for 42 species while *Non-agricultural fire* was recorded for 11. When the category of threat was considered the sub-drivers *Cropping* and *Hunting* affected most of the CR species (39 and 14 species, respectively) and VU species (142 and 49 species, respectively). *Cropping* (114 species) and *Livestock Farming* (41 species) were the sub-drivers affecting most of the EN species (Fig. 4).

Mammal species were most affected for 6 out of the 10 drivers analyzed, ranging from 30% up to 89% of the species depending on the driver, and bird species were the most affected for two drivers (Supplementary Figs. 6-S9). *Agricultural activity* was the top driver for mammals and birds; reptiles accounted for 20 out of the 50 species affected by the driver *Energy Production,* while amphibians were 8 out of the 30 species affected by the driver *Pollution*. No driver was group-specific.

For the driver *Agricultural activity,* 206 species were affected by the sub-driver *Cropping* alone (139 birds, 28 mammals, 27 reptiles, and 12 amphibians); five species were affected by the sub-driver *Livestock Farming* alone (3 birds, 1 reptile and 1 amphibian); and 68 species were simultaneously affected by *Cropping* and *Livestock Farming* (33 mammals, 11 birds, 17 reptiles, and 7 amphibians).

For the driver *System Modification*, 41 species were affected by the sub-driver *Dams* alone (15 birds, 15 reptiles, 6 mammals, and 5 amphibians), 30 species were affected by the sub-driver *Agricultural fire* alone (12 birds; 8 mammals, 8 reptiles, and 2 amphibians); and 9 species by the sub-driver *Non-agricultural fire* alone (5 birds, 3 mammals, and 1 amphibian). For the driver *Overexploitation*, 75 species were affected by the sub-driver *Hunting* alone (38 mammals, 32 birds, and 5 reptiles) and 29 species by sub-driver *Logging and Wood Harvesting* (13 reptiles, 12 birds, 3 amphibians, and 1 mammal).

Mammals were the most affected group for 8 out of the 10 most-frequent sub-drivers, varying from 37% up to 82% of the species depending on the sub-driver. The sub-drivers *Aquaculture, Recreational, Excess Energy, Climate Change and Severe Weather n/i,* and *Droughts* were recorded for mammals only (Supplementary Table 2). Birds were the most affected group for the sub-drivers *Cropping* (155/295 spp. −53%), and *Dams* (17/50 spp. − 34%). Fifteen out of the 37 species (40%) affected by *Mining* were reptiles, and 12 out of 76 species (16%) affected by *Housing* were amphibians.

## Discussion

More than half of the 1172 endangered species in Brazil are being Red-Listed based on the continuing decline in the size, extent and/or quality of their habitats (IUCN criterion B). Moreover, the main driver threatening those species is clear: 2/3 of the 464 vertebrate species analyzed in depth are negatively impacted by agribusiness. Other major threats are related with hunting, urban sprawl, rural settlements, and the construction of hydroelectric dams. Although threats were identified, the availability of national-level quantitative data on population decline (criterion A), on the number of sexually reproductive individuals (criterion C), and estimates of population sizes (criterion D) are highly variable among the Brazilian animal groups – from zero to the majority of them, to reliable quantitative data for some species of mammals and birds – making the use of these criteria to be frequently group-specific.

The causes of deforestation in tropical regions can be direct – i.e., related to land use, and directly affecting the environment and vegetation cover – or indirect – i.e., causes which are related and determine an increase in the demand for actions producing changes in the use of land (Geist & Lambin 2002). We detected that the five most frequent threatening drivers on our analysis are a mix of both direct and indirect causes: among the direct causes are agribusiness, logging, and the implementation of infrastructure such as roads, highways and dams. Indirect threats can be more difficult to identify and measure, ranging from demographic, economic, technological, institutional, and cultural issues (Geist & Lambin 2001). In our analysis, these underlying causes include socio-economic issues, such as population growth and urban sprawl, tourism and industrial activities, and rural settlements, as well as cultural issues, like wildlife hunting, and political issues, like changes in the Brazilian Forest Code (Tollefson 2011; Soares-Filho et al. 2014; Roriz et al. 2017).

Agricultural activities is the second greatest threat to 8000 species threatened with extinction globally, affecting 68% of the species (Maxwell et al., 2016). Our analysis confirms that: agricultural activities affect 47% of the critically endangered species of Brazilian vertebrates, 68% of the endangered, and 73% of the vulnerable species. The conversion of natural habitats to the agriculture is now occurring more rapidly in tropical regions and driven by the demand for commodities such as soybeans, coffee, cocoa, sugar, and palm oil (Curtis et al. 2018). In the case of Brazil, habitat loss is largely driven by deforestation and several studies have indicated both large-scale and slash and burn agriculture as the main drivers. The increase in sugarcane expansion, for example, led to significant changes in land use in the Atlantic Forest (Galindo-Leal & Câmara, 2003), while cattle ranching and soybean cultivation are major drivers for deforestation in Brazilian Amazonia and the Cerrado (Barona et al., 2010; Fearnside 2005; Spring 2018). Agriculture and agribusiness also bring with them several indirect causes, such as the use of fire, recorded for 10% of the species here analyzed. The use of fire in Brazil is closely related to the intensification of agricultural production and opening of pastures, resulting in an increase of fire frequency along the Brazilian agricultural frontier (Carrero & Fearnside 2011; Hantson et al. 2015). Furthermore, agribusiness was tied to 36% of the species for which pollution was identified as a threat, due to the runoff and leaching of agrochemicals or other agricultural residues. This emphasizes the large share of direct and indirect effects agriculture and agribusiness have on threat to species in Brazil.

The opening of roads is a form of infrastructure that has a negative effect on wildlife (e.g. Laurance & Arrea, 2017). Nevertheless, in the next three decades, the total length of additional paved roads could approach 25 million kilometers worldwide (Alamgir et al. 2017). Studies in Brazil estimate that up to 475 million animals may die hit by cars annually, made up of 90% small vertebrates (mainly amphibians), 9% medium-sized vertebrates (such as reptiles and birds) and 1% large vertebrates (such as jaguars and primates)(CBEE, 2018). In our analysis, 10% of the species analyzed experienced threats related to the expansion of the road network, but road kill was identified as a threat to only 2% of them, all mammals, including the maned-wolf *Chrysocyon brachyurus* and the puma *Puma concolor*.

### Underestimating threats

In addition to the effects of deforestation and fragmentation, other drivers seem to be clearly underestimated in Brazil. This is the case for climate change. Climate change can considerably modify the abiotic conditions for the survival of species in the future, increasing the negative effects of habitat loss and fragmentation (e.g. Colombo & Joly, 2010; Mantyka-Pringle et al. 2015; Segan et al. 2016). In fact, a recent meta-analysis to identify the main drivers of global threats have indicated that climate change is mentioned by 40% of the published papers, with an increase of 10% per year (Mazor et al. 2018). Climate projections for the Atlantic Forest in northeastern Brazil point to a temperature rise of around 0.5 °C to 3 °C by the year 2070, and rainfall decrease between 20-25%, whereas projections for the Cerrado point to an increase of up to 3.5 °C and reduced rainfall between 20-35% (PBMC 2015). Scenarios for parts of the Amazonia and the Caatinga are even more worrisome, as those regions may experience above average effects of climate change. The climatic models for the Caatinga – a region whose average rainfall is < 800 mm – indicate an increase of 0.5 °C to 1 °C in the air temperature and a decrease between −10 % and −20 % in the rain during the next three decades (until 2040), with a gradual increase of temperature to 1.5 °C to 2.5 °C and decrease between −25 % and −35% in the rainfall patterns in the period of 2041-2070 (PBMC 2015).

Similar analyses to those presented here indicate that climate change is a major threat to the endangered species worldwide. Climate change appears as a threat to 21% of some 8,000 globally endangered species (Maxwell et al. 2016) and, in the long run, climate change is already recognized as the most troubling threat among birds (BirdLife International 2018). Allek et al. (2018) identified climate change among the most prominent threats for the endangered fauna of Australia, especially in the east of the country, where the largest number of species of amphibians are concentrated. Amphibians are sensitive to climate change (Gascon 2007) and in Australia this is the group with the highest number of Critically Endangered species. Brazil and Australia are both continental-sized countries subject to climate change, harboring rich and endemic faunas. However, while Australia has ca. 230 species of amphibians, Brazil has ca. 1,080 (Segalla et al. 2016). Thus, the expected number of amphibian species threatened by climate change would be certainly bigger in Brazil. Our analysis reveals that climate change is currently listed as a threat to only 2% of the vertebrate species analyzed, and with no amphibians or birds among them. Moreover, there are important synergies between forest fragmentation, climate vulnerability and species threat status (e.g. Jetz et al. 2007; Becker et al. 2016), where fragmented forests tend to be more vulnerable to droughts than intact forests (e.g. Scarano & Ceotto, 2015; Segan et al. 2016). Therefore, the impact of climate change for some specific animal groups in Brazil – like amphibians and birds – is underestimated, likely more pronounced that currently assessed, and aggravated by the advanced state of fragmentation present in several Brazilian terrestrial biomes.

The role exotic species play in threatening species in Brazil also is likely underestimated. Worldwide, the presence of predatory exotic species have caused numerous species extinctions, with the best studied impacts being those of cats, rats and dogs (Jones et al., 2008; Doherty et al., 2016). Of the 233 bird species here analyzed, invasive or exotic species were recorded as a threat for only 15. Of these, 60% were related to predation of nests by rats and mice (3 CR, 3 EN and 3 VU). Dogs are already recognized as a conservation problem in the Atlantic Forest, becoming the most frequent recorded species among all mammal locally (Srbek-Araujo & Chiarello, 2008; Paschoal et al. 2012; Lessa et al. 2016). However, in our analysis, the negative impact of dogs was rarely reported as a threat.

Eleven percent of the species we analyzed were impacted by threats associated with water management. Among these, for 9 out of 10 species the implementation of hydroelectric dams was the main threat. In Brazil, large hydro dams are mainly located and planned for the Amazon, generating debate on their negative impacts, including the displacement of human populations due to the flooding of indigenous territories and habitat loss for vertebrates. There is also debate on whether the energy they produce is actually green (Benchimol & Peres, 2015; Lees et al., 2016). In our analysis, two of the species affected by hydro dams are the Amazon river-dolphin *Inia geoffrensis* (EN), and the recently-described Brazilian species *Inia araguaiensis* (Hrbek et al., 2014), found in the Araguaia River basin. This species is on the process of being Red-Listed (Araújo & Wang 2015). Also, in spite of previous and new evidence (Carter & Rosas 1997; Palmeirim et al. 2014; Groenendijk et al. 2015) hydroelectric dams were not identified as a threat to the giant river otter *Pteronura brasilensis* (VU) in the official documents we analyzed. Large hydroelectric reservoirs often greatly increase the extent of freshwater environments, but these often provide poor quality habitats for aquatic biota (Palmeirim et al., 2014). Large dams also profoundly alter the structure of terrestrial biota with species isolated on the islands formed (Benchimol & Peres, 2015; Lees et al., 2016). In the current scenario where 2,215 hydro dams are planned for Amazonia (Finer & Jenkins 2012; Tundisi et al. 2014; Anderson et al. 2018), the conservation consequences are worrisome. Given the high species richness and endemism of Amazon region, and considering that Brazil hosts both most of the impacted area and most of the projected hydro dams, the threat those structures poses to the regional fauna seem underestimated and their impact must be correctly assessed.

### Synergies between threats

Biodiversity loss may be intensified in response to additive, synergistic or antagonistic effects (Pereira et al., 2010; Maxwell et al., 2016). The synergy between threats can worsen the situation of species already threatened (e.g. BirdLife International 2018) and this may occur because the combined effect of two threats may be greater than the additive effect of these threats separately (Allek et al., 2018). Identifying these synergies is important both to quantify the risk of extinction and to prioritize threat mitigation (Ducatez & Sjine, 2017; Allek et al., 2018). Recognizing synergies and trade-offs in a resource-constrained scenario, with a focus on different targets, can minimize efforts and optimize spending on conservation (Di Marco et al., 2015).

However, few studies address the role of multiple threatening drivers (Ducatez & Sjine, 2017; Mazor et al., 2018). In our analysis, the groups with the most threatened species (birds and mammals) were also the groups with the most co-occurrence of threats. The most frequent sub-driver (*Cropping*) was recorded for species of all categories of threat in all the animal groups here analyzed. Cropping was recorded in association with two of the other three sub-drivers of Agricultural Activities (*Livestock* and *Timber Plantations*) with its association with *Livestock Farming* accounted for 22% of the species affected by Agricultural Activities. Integrated predictive studies (e.g. Symes et al. 2018) on the relative and synergistic effects of and threats on the Brazilian biodiversity are still scarce (e.g. Gouveia et al. 2016). Such detailed analyses will be hampered if some threats are underestimated.

### Criterion B, the lack of basic information and the need for refined evaluations

Gaps in the basic knowledge on the distribution of some Brazilian species may compromise their conservation (e.g. Bernard et al., 2011; Sousa-Baena et al. 2013; Oliveira et al. 2017). Some species indeed have large extent of occurrences (hereafter EOO, the estimate of dispersion of risk) built based on a few scattered records. However, a large EOO based on isolated and broadly spaced points may underestimate the real situation of those species, especially for those where the EOO is overall under strong pressure. In this situation, data that would allow for estimation of the area of occupancy (hereafter AOO, or the actual best estimate of distribution) would help to reveal this fragmention (e.g. Jetz et al., 2008). Accurate data to allow for the estimation of AOO is these situation is a high research priority. On the other hand, presenting artificially smaller EOO and AOO as a result of poor data will result in higher threat status and overestimate risk (IUCN, 2012). Most of the Brazilian species classified under criterion B based on their EOO (B1) are mammals, reptiles, amphibians, land invertebrates and aquatic invertebrates; and based on their AOO (B2), birds and freshwater fish. The prevalence of criterion B and the rare use of criterion E are observed worldwide (e.g. Collen et al. 2016). But, in an utopian scenario where there was no such high loss or degradation of habitats in the country, many of the Brazilian species would actually be classified as “Data Deficient” due to the total lack of basic data on their population declines (criterion A), on the number of sexually reproductive individuals (criterion C), and on estimates of population sizes (criterion D). Currently, 1,670 of 12 556 species evaluated in Brazil are “Data Deficient” (ICMBio, 2018). The higher use of criterion B exposes, in fact, a worrisome combination of severe and fast habitat loss and the lack of the most basic population information for most of the endangered species in Brazil, a situation that must be reversed.

Part of this problem can be reduced with regional assessment initiatives. In fact, IUCN does support and encourages regional Red Lists (IUCN 2012) and in a country with continental dimensions such as Brazil the production of more refined state lists may fill some of the knowledge gaps necessary to better classify some species. In the case of smaller states and for those with more data and established technical expertise, regional assessments could be a better alternative. These regional Red Lists can better identify threatened populations or those more likely to decline on a more detailed spatial scale (e.g. De la Torre et al. 2018), allowing the development of a strategy to prevent local population declines that eventually lead an entire species to become threatened with extinction on a wider level. However, currently, only eight of the 27 Brazilian states (Bahia, Espírito Santo, Minas Gerais, Pará, Paraná, Rio de Janeiro, Rio Grande do Sul and São Paulo) have their state Red Lists. Similar initiatives must be encouraged for the other Brazilian states, especially considering that in 2011 legal responsibility for surveillance and enforcement of administrative penalties involving flora, fauna and environmental licensing was transferred from the federal agency (IBAMA) to state and municipal environmental agencies. Therefore, in this case, state and regional Red Lists would have practical consequences.

### A pessimistic conservation scenario ahead

As of November 2018, Brazil elected a new president, aligned with a far-right agenda, identified by several analysts as very detrimental to the future of the country’s environment and biodiversity, especially to the Amazon and indigenous and traditional peoples, and also resulting in the destabilization of the global climate (e.g. Carneiro Filho, 2018; Fearnside & Schiffman 2018). As soon as he took office, the president-elect reduced the role of Brazil’s environmental ministry and the environmental agencies IBAMA (surveillance and environmental licensing) and ICMBio (protected areas and biodiversity management) (e.g. Abessa et al. 2019; Phillips 2019). His appointed Minister of the Environment public declared he was favorable to freeze the creation of new protected areas and Indigenous lands, plus his intention to “analyze in detail” – including the possibility to degazette – the entire 334 federal protected areas in Brazil (e.g. Kaiser 2019; Borges & Branford, 2019). The minister also publicly declared to be favorable to open protected areas to mining, reduce licensing requirements for major infrastructure projects such as dams, industrial waterways, roads and railways (Branford & Borges, 2019; Gonzales, 2019). The president-elect was deeply supported by the most outdated and least environmental friendly part of Brazil’s agribusiness, industrial and commercial sectors. Considering that the main threats identified in our study are directly related to the agribusiness, mining and infrastructure sectors – the basis of the country’s economy – such combination of political and economic factors projects a pessimistic conservation scenario ahead for Brazil’s biodiversity. Under the current role played by the president-elect and his Minister of the Environment the number of threatened species in Brazil is poised to increase and the degree of threat of those already red-listed will definitively worse in the near future.

## Author contributions

Study design: EB; data analysis: EB, FSB, VEFF; writing the article: EB, FSB, TEL Jr.

## Acknowledgements

We thank Ednaldo Monteiro da Silva for handling and sorting part of the data. This manuscript is partially based on FSB’s Honors Thesis to receive her B.Sc. in Biology/Environmental Sciences at UFPE, and we would like to thank Drs. FPL Melo and JP Souza-Alves for taking part in her evaluation committee. EB would like to thank the Brazilian Conselho Nacional de Desenvolvimento Científico e Tecnológico - CNPq for a fellow grant and the Departament of Zoology at UFPE for supporting his research on biodiversity conservation in Brazil.

## Ethical Standards

The research here presented did not involve human subjects, experimentation with animals and/or collection of specimens.

