## Supplementary Information for "What threatens Brazilian endangered species and how they are Red-Listed"

FERNANDA SILVA DE BARROS - Laboratório de Ciência Aplicada à Conservação da Biodiversidade, Departamento de Zoologia, Universidade Federal de Pernambuco, Recife, Brazil

VITÓRIA EDNA FERNANDES FELIX - Laboratório de Ciência Aplicada à Conservação da Biodiversidade, Departamento de Zoologia, Universidade Federal de Pernambuco, Recife, Brazil

THOMAS E. LACHER JR - Biodiversity Assessment & Monitoring Lab, Deptartment of Wildlife and Fisheries Sciences, Texas A&M University, College Station, Texas, USA

### Methods

Five modifications were necessary in the categories proposed by Salafsky et al. (2008), and all were made with the intention of increasing the clarity and adequacy of the categories originally proposed:

1- Inclusion of the sub driver *Pollution - unknown* in the driver Pollution so we could accommodate threats related to pollution but whose causes were not clearly defined.

2- Inclusion of the sub driver *Climate change - unknown* in the driver Climate Change so we could accommodate threats related to climate change but whose causes were not clearly defined.

3- In the classification proposed by Salafsky et al. (2008), the sub driver *Fire and fire suppression* is included in the driver Natural System Modifications and is defined as "fire suppression to protect homes, inappropriate fire management, escaped agricultural fires, arson, campfires,

fires for hunting”. However, in Brazil most forest fires are related to the clearing and preparation of areas for agriculture and pasture (both small and large scale) (Hantson et al. 2015; Andela et al. 2017; da Silva et al. 2018). Other uses of fires are rare and we therefore divided the sub-driver *Fire and fire suppression* in two: 1) *Agricultural fire*, to accommodate all threats related with the use of fire for opening new or cleaning existing crop and/or pastures areas, and for situations where fire is used as part of the harvesting procedures, like in the case of sugarcane; and 2) *Non-agricultural fire*, to accommodate threats related with arson, campfires, or cases where fire was mentioned as a threated without any further details. Thus, for clarity, these sub-drivers are presented separately.

4- Considering that in Brazil clearing for crops and pasture is recognized as the most important driver of deforestation and loss of original vegetation cover (e.g. Veríssimo et al., 1996; Arima et al. 2005; Carrero & Fearnside 2011; Barreto et al. 2013) all threats described as habitat loss, habitat fragmentation and decline of habitat quality and/or area of occurrence were credited under the driver Agriculture and aquaculture, sub driver *Cropping*.

5- We included in the driver System Modification, sub driver *Others,* all threats attributed to the recent changes in the Brazilian legislation (namely the Forest Code and the cave protection law (e.g. Galetti et al. 2010; Stan et al. 2015; Leal et al. 2017), or those pointing to problems due to area overlap between Indigenous Lands and protected areas, and those related to problems due to illegal land grabs.

6- Most threats related to diseases or pathogens did not specify if those were native or exotic. Therefore, in these cases they were credited to the driver Invasive and other problematic species and genes, sub driver *Invasive non-native/alien species*.

**Supplementary Figures**

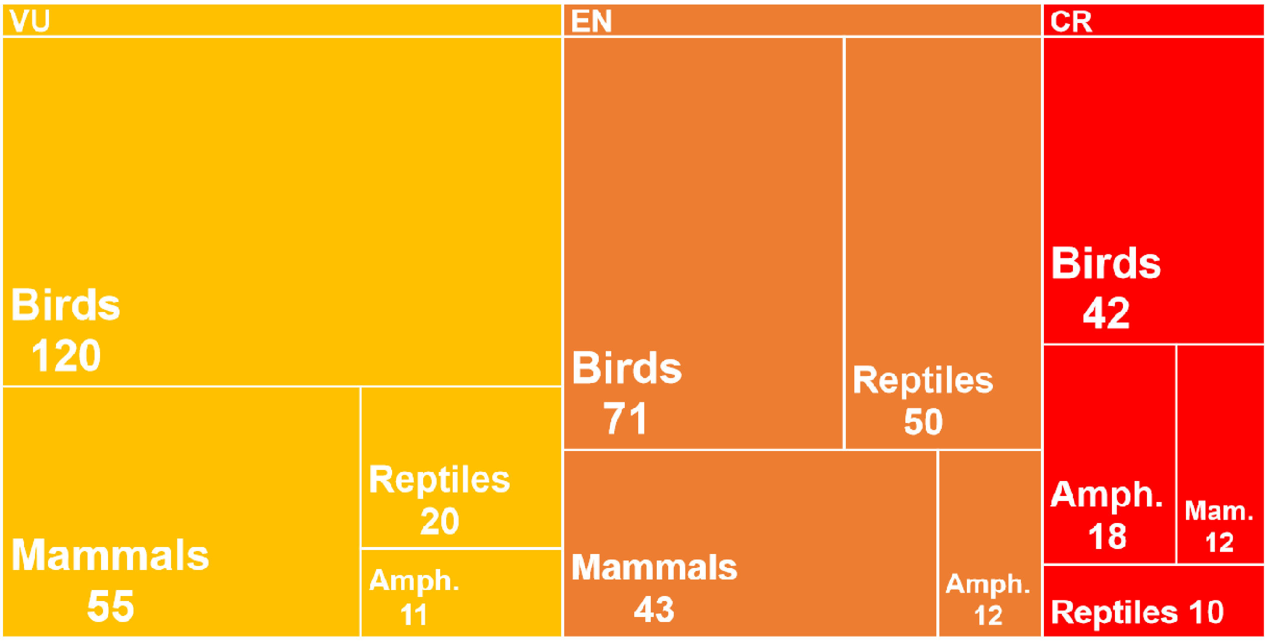


Supplementary Figure 1: Conservation status of 464 species of birds, mammals, reptiles and amphibians officially listed as threatened in Brazil. VU = Vulnerable; EN = Endangered; CR = Critically Endangered.

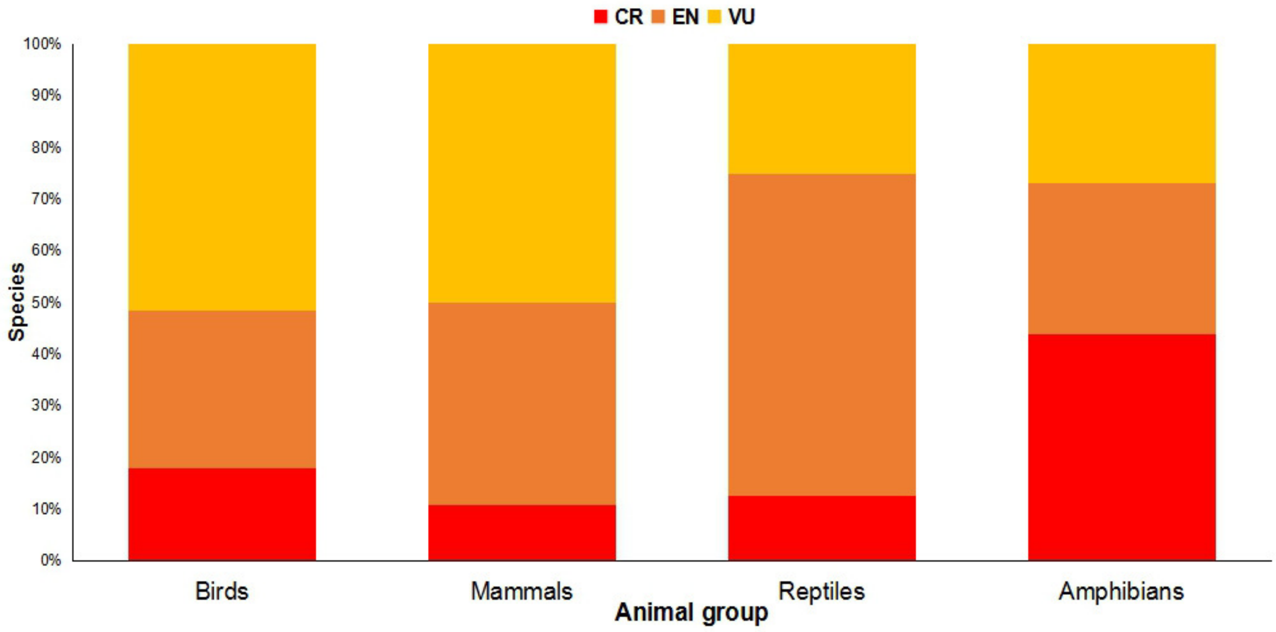


Supplementary Figure 2. Conservation status (in percentage) of 464 species of terrestrial vertebrates officially listed as threatened in Brazil. CR = Critically Endangered; EN = Endangered; VU = Vulnerable. Birds, n = 233 species; mammals, n = 110 species; reptiles, n = 80 species; and amphibians, n = 41 species.

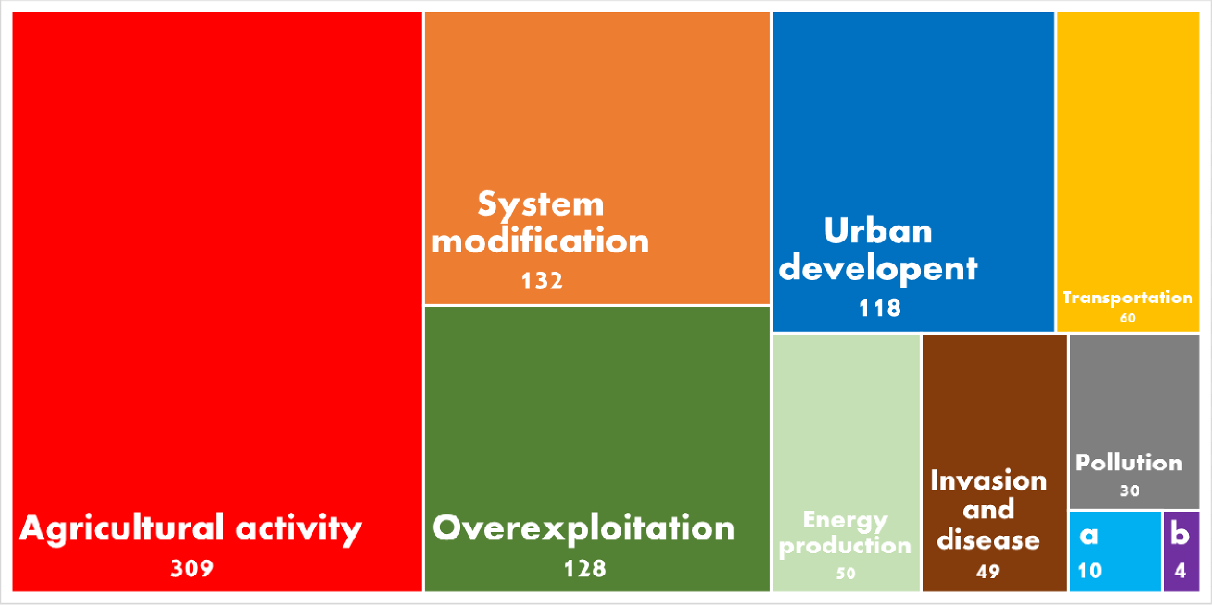


Supplementary Figure 3: Top-10 threatening drivers which affect the conservation status of 464 species of terrestrial vertebrates officially threatened in Brazil. Drivers were classified according to Salafsky et al. (2008). Numbers represent the frequency each driver was recorded. a, Climate change; b, Human disturbance.

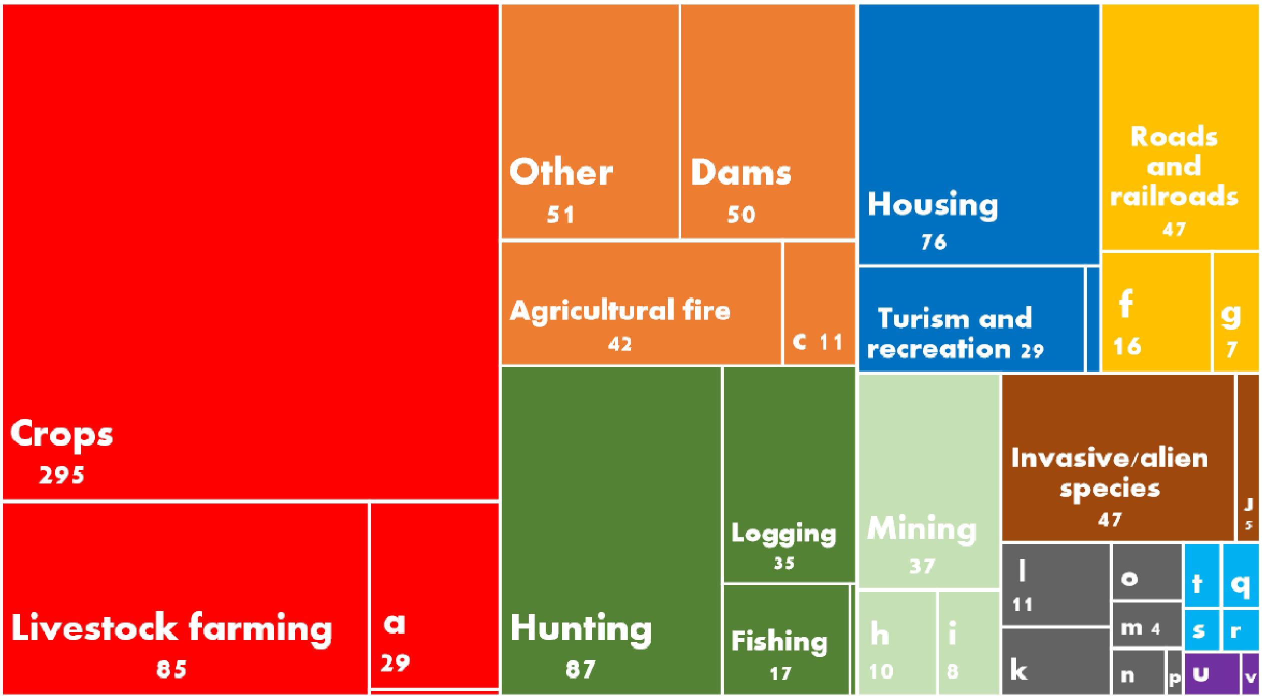


Supplementary Figure 4: Sub-drivers affecting the conservation status of 464 species of terrestrial vertebrates officially threatened in Brazil. Drivers and sub-drivers were classified according to Salafsky et al. (2008). Sub-drivers with the same color belong to the same driver, as in Figure S3. Numbers represent the frequency each sub-driver was recorded. a, Timber plantation; b, Aquaculture; c, Non-agricultural fire; d, gathering plants; e, Industrial; f, Service lines; g, Shipping lanes; h, Renewable energy; i, Oil and gas; j, Problematic native species; k, Pollution n/i; l, Agricultural; m, Domestic waste; n, Industrial; o, Air-bone; p, Excess energy; q, Storms and flooding; r, Extreme temperature; s, Climate change n/i; t, Droughts; u, Recreational; v, War, civil unrest and military exercises.

| Ranking | Animal group | | | |
|---|---|---|---|---|
| | Mammals | Birds | Reptiles | Amphibians |
| 1 | Crops | Crops | Crops | Crops |
| 2 | Hunting | Hunting | Housing | Housing |
| 3 | Other | Livestock farming | Livestock farming | Livestock farming |
| 4 | Livestock farming | Dams | Dams | Other |
| 5 | Roads and railroads | Invasive/alien species | Mining | Dams |

Supplementary Figure 5: Most important conservation sub-drivers affecting 464 species of terrestrial vertebrates officially listed as threatened in Brazil and their respective rankings according to animal group. Drivers were classified according to Salafsky et al. (2008)

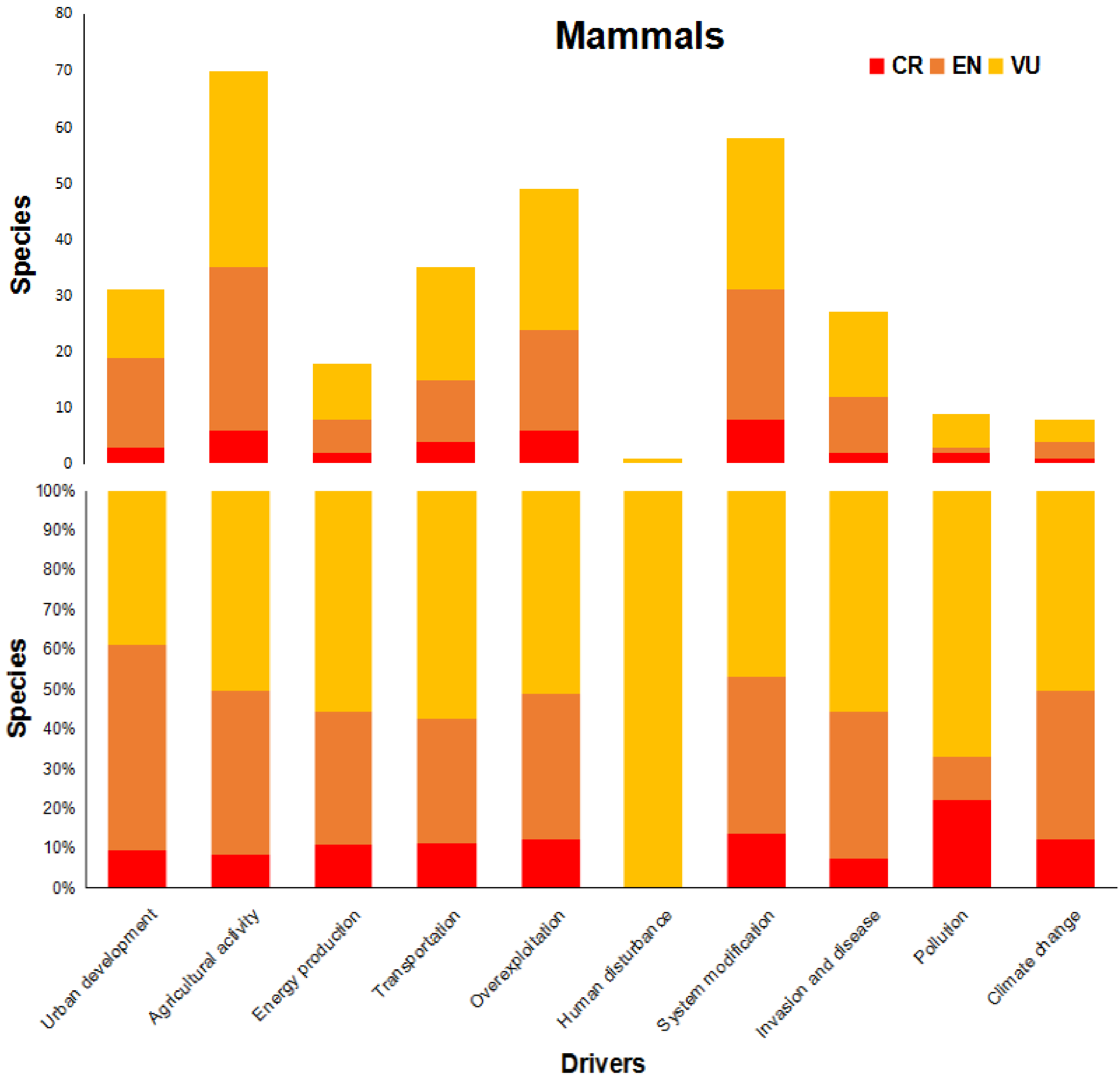


Supplementary Figure 6: Threatening drivers which affect 110 species of threatened mammals in Brazil. Drivers were classified according to Salafsky et al. (2008). CR = Critically Endangered; EN = Endangered; VU = Vulnerable.

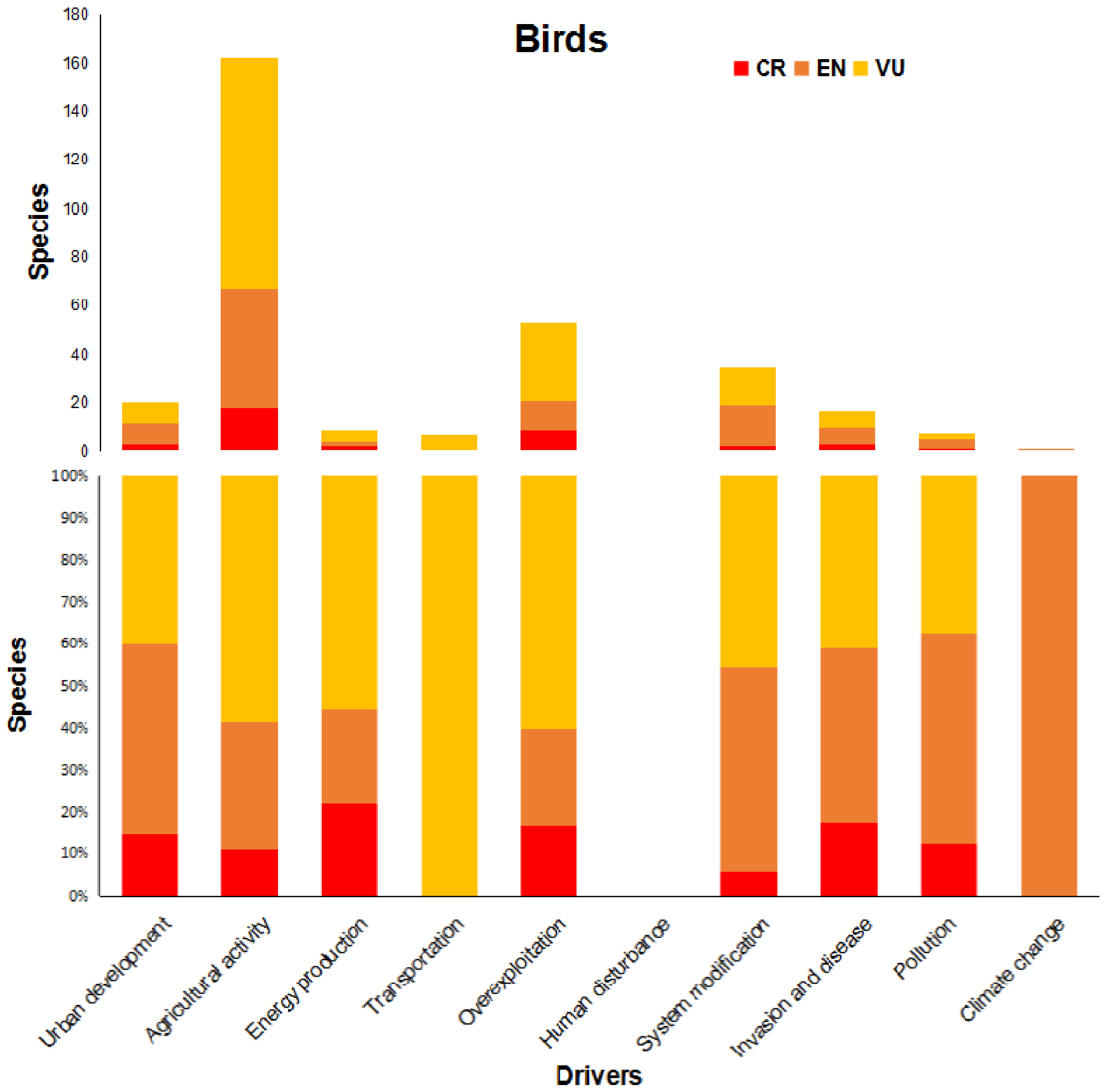


Supplementary Figure 7: Threatening drivers which affect 233 species of threatened birds in Brazil. Drivers were classified according to Salafsky et al. (2008). CR = Critically Endangered; EN = Endangered; VU = Vulnerable.

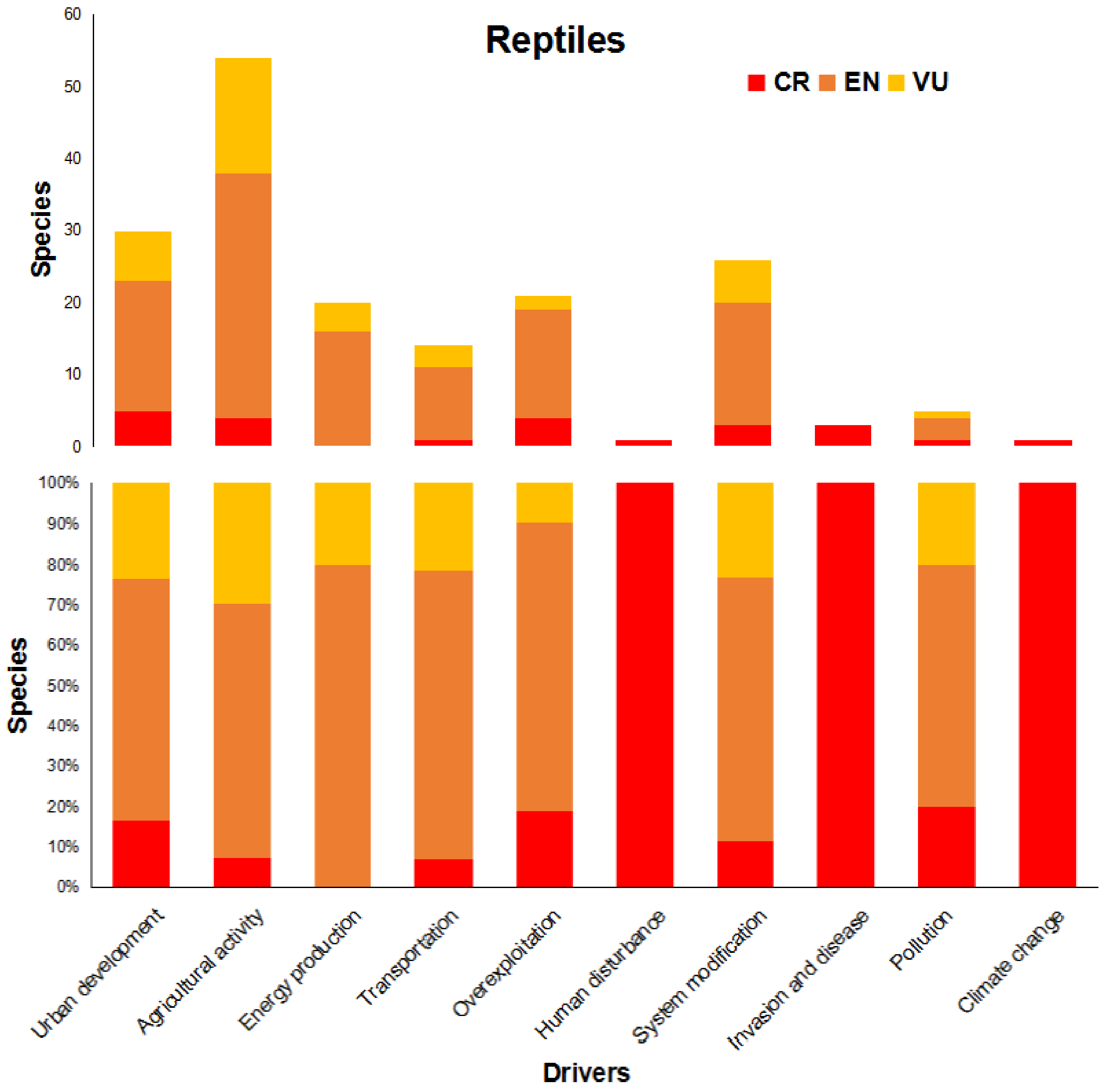


Supplementary Figure 8: Threatening drivers which affect 80 species of threatened reptiles in Brazil. Drivers were classified according to Salafsky et al. (2008). CR = Critically Endangered; EN = Endangered; VU = Vulnerable.

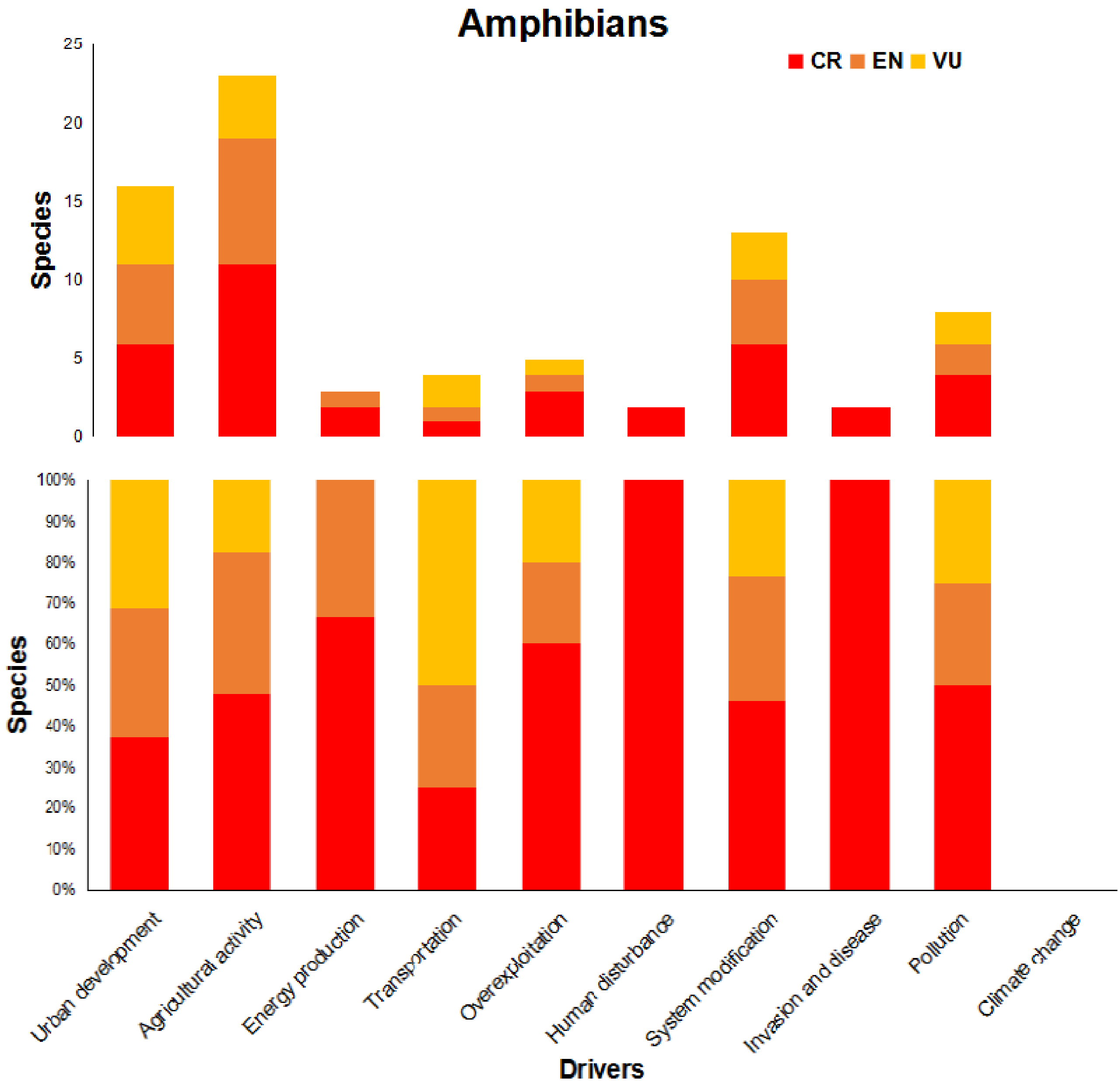


Supplementary Figure 9: Threatening drivers which affect 41 species of threatened amphibians in Brazil. Drivers were classified according to Salafsky et al. (2008). CR = Critically Endangered; EN = Endangered; VU = Vulnerable.
